# *UNC5H3* expression during early development in the chicken is associated with populations at the dorsal apex of the neural tube

**DOI:** 10.1101/2025.02.28.640710

**Authors:** Ruth Klafke, Natalie Harriman, Richard J.T. Wingate, Andrea Wizenmann

## Abstract

The Netrin receptor Uncoordinated-5 receptor C (UNC5C) or UNC5HA has been shown to regulate cell migration in cerebellum and cortex, to control the guidance of the axons in the developing corticospinal tract and to act as a dependence receptor in apoptosis in mouse and rat. We have examined the distribution of *UNC5H3* using whole-mount mRNA *in situ* hybridisation in the embryonic chick, concentrating on its early expression in the mesencephalic/metencephalic region in relation to known ligands, *NETRIN1* and *NETRIN2*. From E2 (embryonic day 2), while the latter are confined to the ventral midline, *UNC5H3* is expressed exclusively at the dorsal midline from the diencephalon caudally. Later in development, *UNC5H3* is maintained in the rhombic lip (E5 onwards) and subsequently expressed in its putative derivatives; cerebellar granule cells and nuclei within the avian auditory hindbrain complex and the inferior and superior olive.

## 1. Results and discussion

Axon navigation and migration of neurons depends on attractive and repulsive guidance cues during the development of the central nervous system (CNS). Netrin proteins provide a small phylogenetically conserved family of secreted proteins important for guiding axonal growth cones to their targets (Serafini et al., 1994; (Bloch-Gallego et al., 1999). Netrins are bifunctional: they attract some axons and repel others (Alcantara et al., 2000). Netrin receptors of the deleted in colorectal carcinoma (DCC) family mediate growth cone attraction (Keino-Masu et al., 1996), while those of the Unc5 family mediate repulsion (Tessier-Lavigne and Goodman, 1996; Hong et al., 1999). In chick the sequences of four *NETRIN* genes are found on pub med. Two, *NETRIN-1* and *NETRIN-2*, have been described in chicken (Serafini et al., 1994), while little is known of the expression of their likely receptors, which include the three Unc5 family members. All three UNC5 proteins bind NETRIN-1 (Leonardo et la. 1997). Mutations of these three genes in mice show caudal midbrain and rostral hindbrain abnormalities (lane et al., 1992, Ackermann et al., 1997). Unc5 binding to GPC3 (glypican-3) guides pyramidal neurones in the mouse cortex (Akkermans et al., 2022). We have hence examined *UNC5H3 (also called UNC5C)* expression in relation to *NETRIN-1* and *NETRIN-2* expression in the midbrain and rostral hindbrain to evaluate if the expression is similar to that in mouse.

Within the developing CNS strong *UNC5H3* expression is detected as early as HH8 (Hamburger and Hamilton stage 8; Hamburger and Hamilton, 1951) in the area of the prospective head (inset in Fig. 1A). Transverse sections (section plane marked by line in inset of Fig. 1A) of this stage revealed that the expression is restricted to the head mesenchym and absent in the neural ectoderm (Fig. 1A). After the neural tube has closed, *UNC5H3* is strongly expressed in the roof plate of the mesencephalon from stage 10 onwards to HH24 (Fig. 1B, 2, and 4). This is consistent with the expression found in the developing mouse embryo, where high levels of *Unc5h3* expression is detected in the roof of the mesencephalon, diencephalon, cerebellar plate, and lateral regions of the fourth ventricle at embryonic day (E) 10.5 (Przyborski et al., 1998). A similar expression is found in zebrafish (Yang et la., 2013). Although in mouse the dorsal expression domain of *Unc5h3* is much broader than the restricted roof plate expression found in the chicken embryo. Nevertheless, there seems to be a conserved function of Unc5h3 in the dorsal part of the brain throughout the species. Interestingly, *UNC5H3* is not expressed in neurons born adjacent to the roof plate of the midbrain such as those of the mesencephalic trigeminal nucleus, whose axons pioneer the descending lateral longitudinal tract receptor (Chedotal et al., 1995; Hunter et al., 2001). A whole-mount in situ hybridisation at HH20 showed that *UNC5H3* is also expressed in the cranial ganglia VII and VIII, branchial arches and ventral head mesenchym (Fig. 1C arrowhead), while central neural expression is still confined to the roof plate of the diencephalon, mesencephalon and rhombic lip (Fig. 1C arrows, and D).

**Figure 1.**
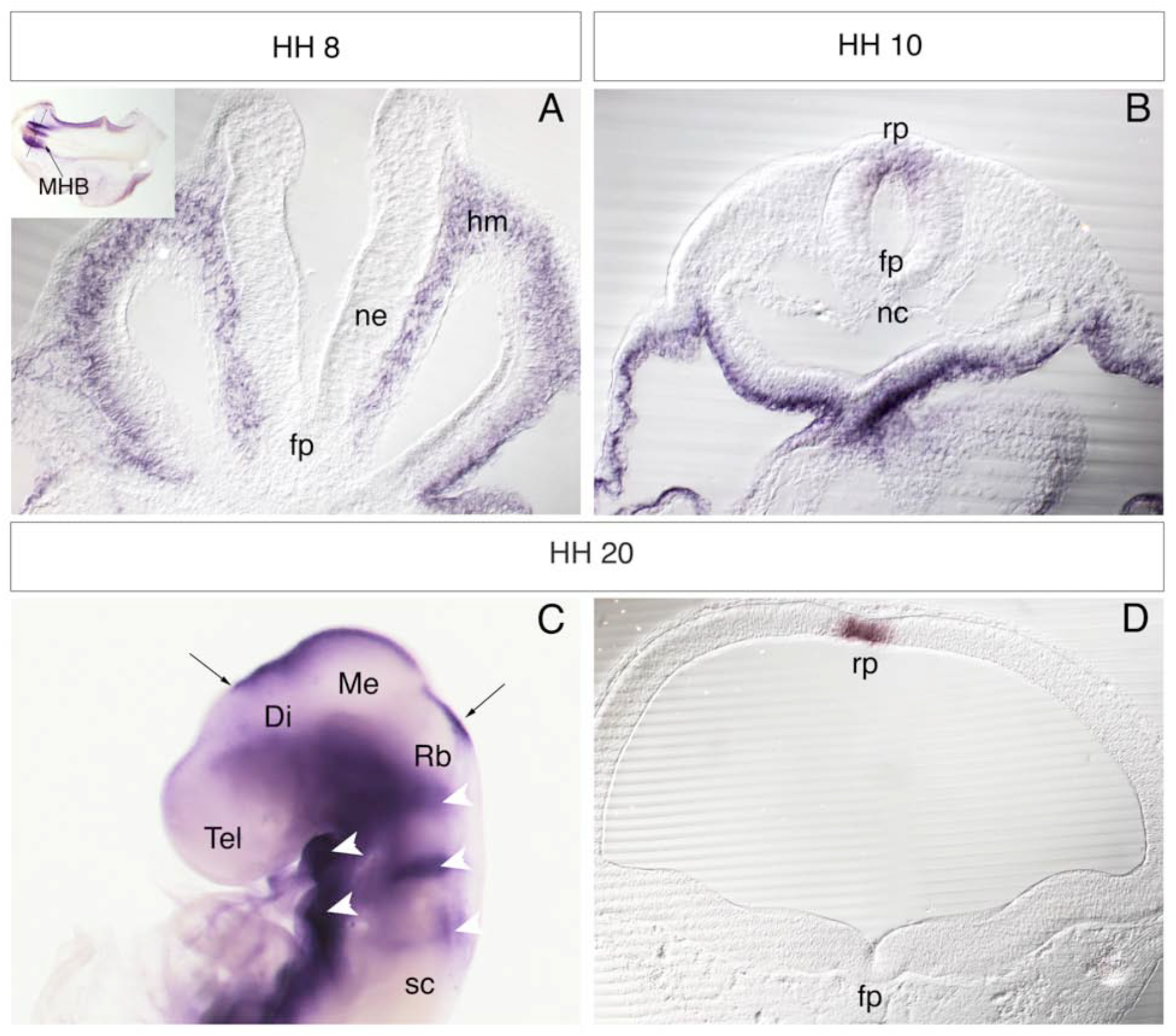
Whole-mount mRNA *in situ* hybridisation of *UNC5H3* on different stages of development. In (A), (B), and (E) dorsal is up and ventral is down. (A) Transversal sections through the presumptive head region at stage 8 shows strong expression in the mesenchyme up to the mid-/hindbrain boundary (see whole mount staining in insert as well as section plane). (B) The earliest expression of *UNC5H3* in the roof plate can be seen at HH10. This roof plate expression continous up to stage 20. (C - D). At stage 20 *UNC5H3* expression in the brain is restricted to the roof plate of the mesencephalon (E) to the synencephalon (arrows), in the rhombencephalon (rhombic lip), the branchial arches and the otic vesicle. (Di - diencephalon; fp - floor plate; hm - head mesenchym; Me - mesencephalon; MHB - mid-hindbrain-boundary; nc - notochord; ne - neural epithel; ov - otic vesicle, Rb - rhombencephalon; sc - spinal cord; so - somite; Tel - telencephalon)

To study the temporal and spatial relation between the expression pattern of *UNC5H3* and two of its ligands *NETRIN1* and *NETRIN2* we performed double whole-mount *in situ* hybridisation at different stages with antisense probes for either *UNC5H3* and *NETRIN-1* or *UNC5H3* and *NETRIN-2*. At HH12 a strong *UNC5H3* expresssion is detected in the roof plate of the midbrain and spinal cord (Fig. 2A, B, G), whereas *NETRIN1* is expressed in the basal plate of the midbrain (Fig. 2A) and floor plate of the spinal cord (Fig. 2B). The *NETRIN-2* expression is also seen in the ventral part of the midbrain, but the expression domain includes basal and alar plate (Fig. 2G). When development proceeds transcripts of *UNC5H3* could still be detected in the roof plate of the midbrain and spinal cord of HH17 embryos (Fig. 2C, D, H, I). Expression of *NETRIN-1* and *NETRIN-2* is confirmed to the ventral part of the midbrain with *NETRIN-2* showing an expression domain expanding more dorsally (Fig. 2C, H). In the spinal cord *NETRIN-1* expression is restricted to the floor plate whereas *NETRIN-2* transcripts are absent from the ventral midline and present in two stripes in the ventro-medial part (Fig. 2D, I). The same situation is found in embryos three stages later ar HH20 (Fig. 2E, F, J, K), now with an additionally slight expression of *NETRIN-1* in the medial part of the spinal cord (Fig. 2F) and weak *UNC5H3* expression in the overlaying ectoderm of the roof plate (Fig. 2F, K).

**Figure 2.**
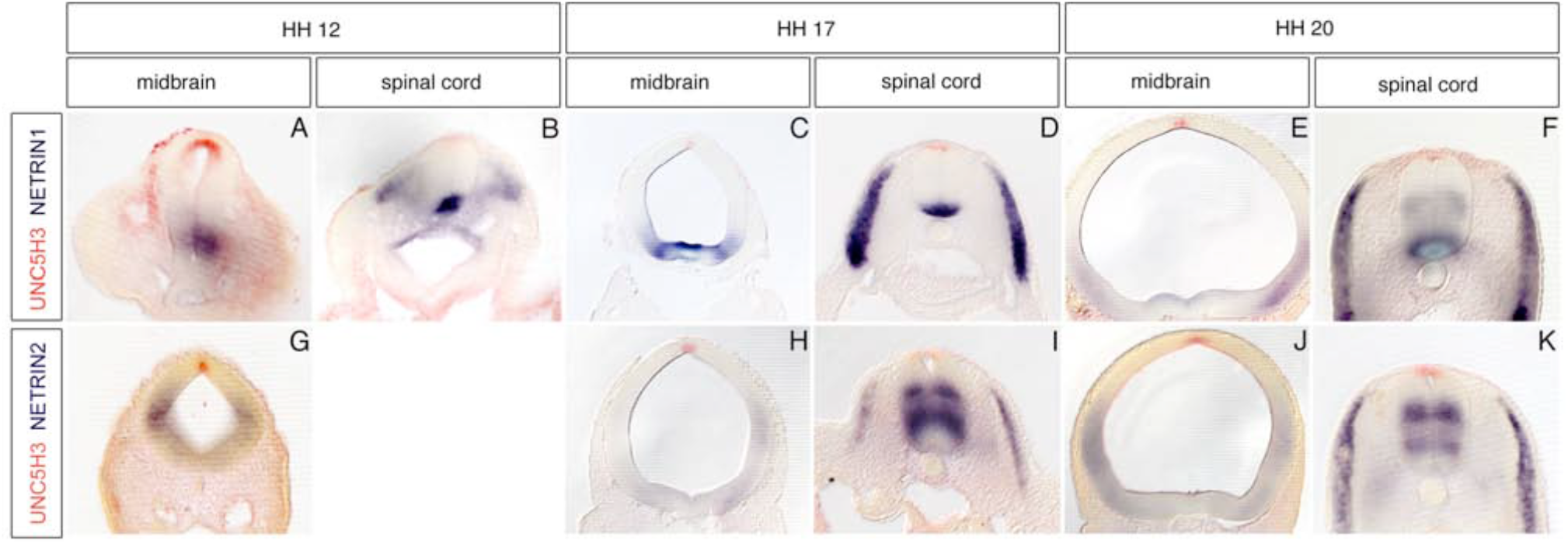
Transverse sections of mRNA *in situ* hybridisation of *UNC5H3* (red) together with *NETRIN 1* (A-F, blue) or *NETRIN 2* (G-K, blue) at different stages of development. Dorsal is up and ventral is down. (A-K) The expression pattern of *UNC5H3* (blue) is non overlapping and distinct from that of *Netrin-1* and *Netrin 2* (red) in the midbrain and in the spinal cord. In both midbrain (A, C, E, G, H, J) and spinal cord (B, D, F, I, K) *UNC5H3* is expressed in the dorsal roof plate and *NETRIN 1* and *NETRIN* 2 in the basal plate. At stage 17 *NETRIN 1* and *NETRIN 2* are in addition expressed in the somites (dermamyotome) (D,I). (F) At stage 20 *NETRIN 1* expression in the spinal cord has expanded more medially.

Whole-mount in situ hybridisation on HH16 with *UNC5H3* combined with immunostaining for the motor neuron-specific proteins, ISLET-1/2 (Ericson et al., 1996), show that neurons expressing ISLET-12/ (Fig. 3B,D) in ventral spinal cord and midbrain and in dorsal midbrain do not express *UNC5H3* (Fig. 3A,C). *UNC5H3* is only expressed in the roof plate at HH16. An *in situ* hybridisation for *UNC5H3* on paraffin sections at HH24 still shows a clear expression domain of *UNC5H3* in the roof plate of the spinal cord (Fig. 4A) and midbrain (Fig. 4B). High levels of expression are now also detected in the dorsal mesenchym and low levels are seen in the ventral motoneuron domain of the spinal cord and the dorsal root ganglia (Fig. 4A). Consecutive sections hybridised with a probe for *ISLET-1* (Ericson et al., 1992; Fig. 4C, D), show a clear overlap in the expression of the two genes in the motorneuron domain of the spinal cord and the dorsal root ganglia (Fig. 4A, C, E). In the midbrain in addition to the roof plate expression there are now two ventral *UNC5H3*-positive domains observable (Fig. 4B). These expression domains could be considered to be the mesencephalic oculomotorius nuclei by the coexpression of *ISLET-1* within (Fedtsova and Turner, 2001; Fig. 3D, F).

**Figure 3.**
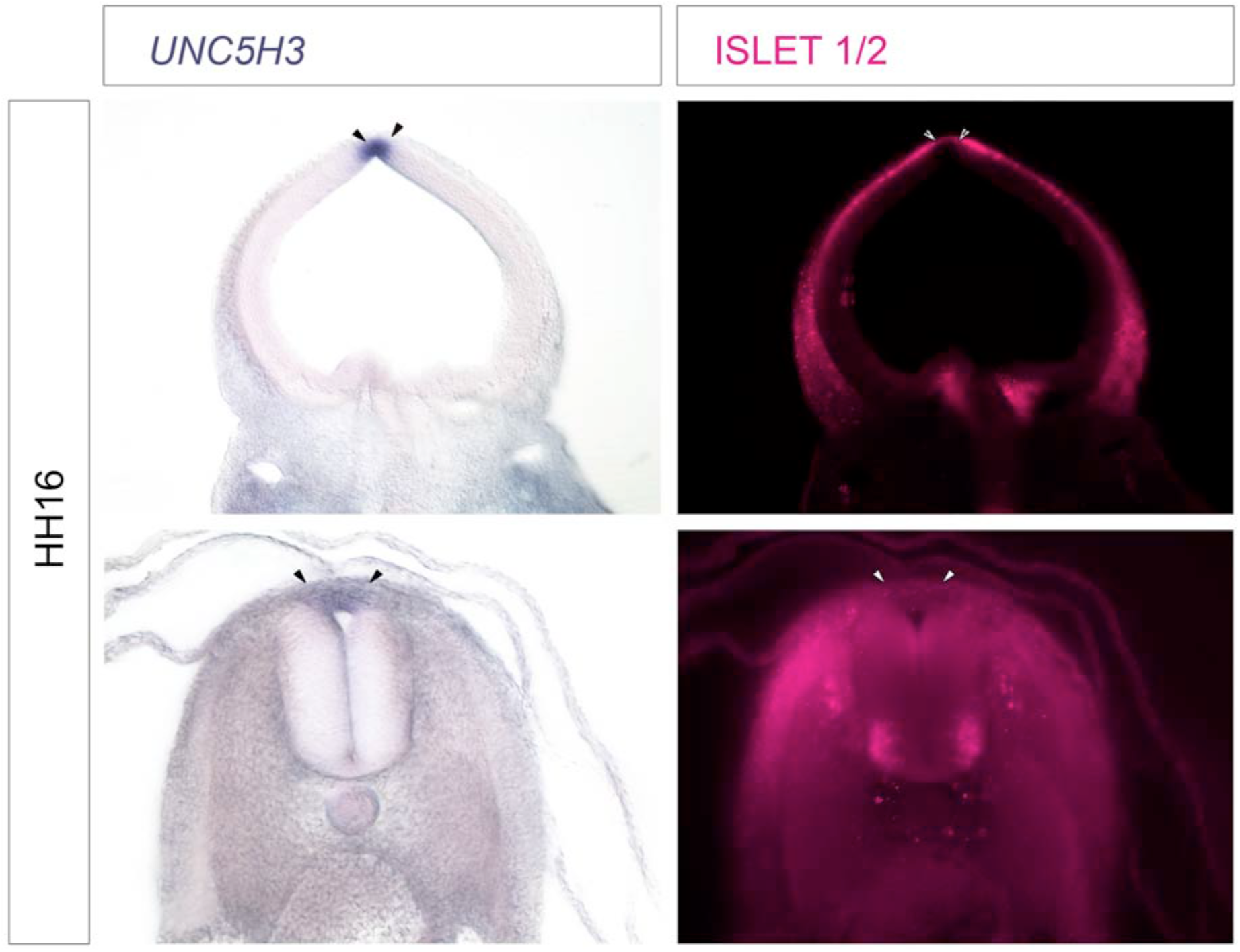
Transversal sections of the mesencephalon and spinal cord at HH16. In all pictures dorsal is up, ventral is down. (A-D) Double labeling with an antibody against ISLET-1/2 shows that *UNC5H3* is not expressed in ISLET-1/2 positive cells in the dorsal midbrain (A-B) or dorsal spinal cord (C-D). The roof plate extent is indicated by arrowheads.

**Figure 4.**
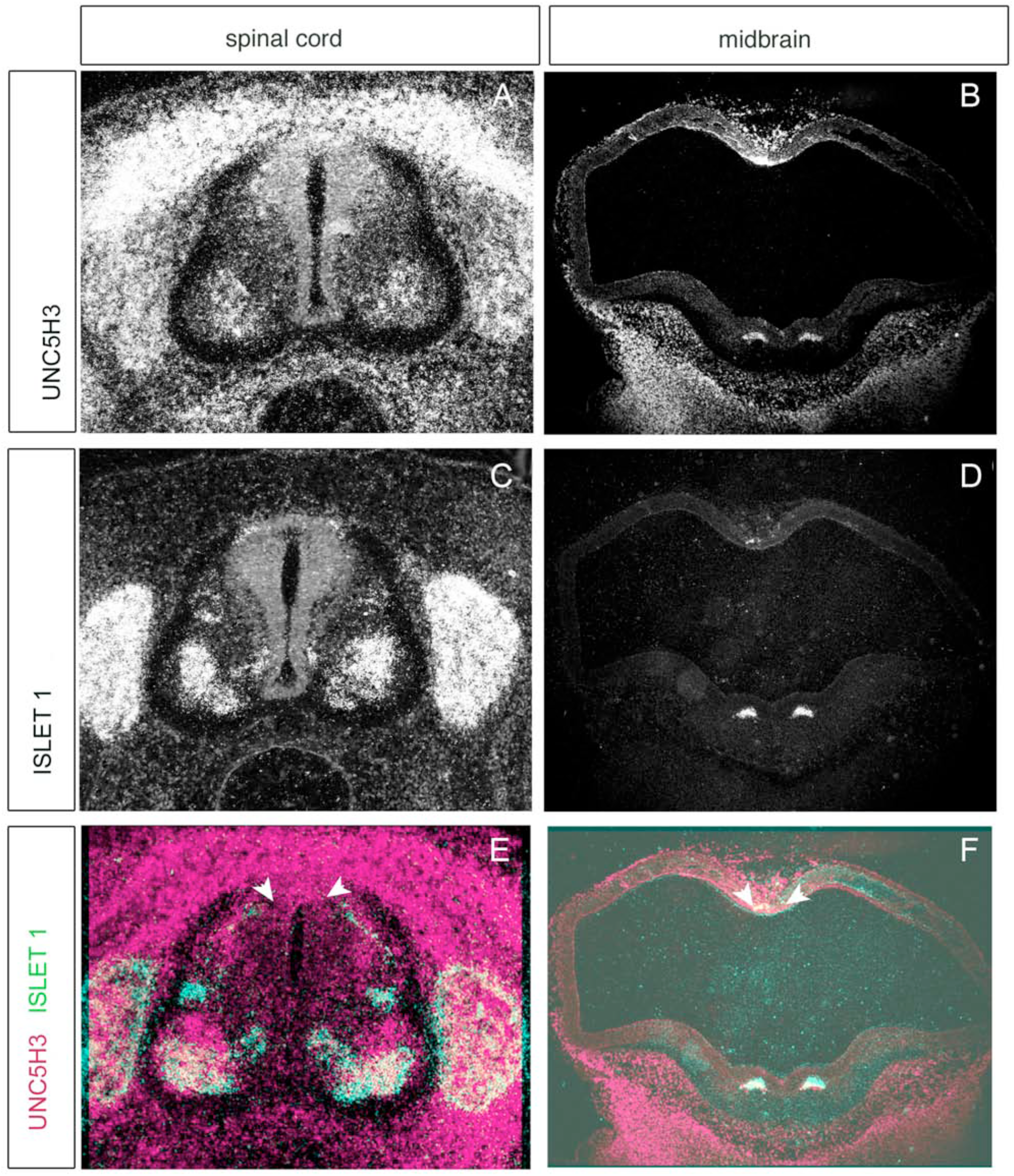
Whole-mount *in situ* hybridisation of *UNC5H3* and *ISLET 1* in transversal section of midbrain and spinal cord at stage 24. Dorsal is up and ventral is down. (A) and (B) show *UNC5H3* expression in spinal cord and midbrain, respectively. (C) and (D) display the *ISLET-1* expression in these regions. The overlay of (A) with (C), and (B) with (D) is shown in (E), (F), respectively. At HH24 *UNC5H3* (red) is co-expressed with *ISLET-1* in motoneurons (green) in the spinal cord (E) and ventral midbrain cells (F) (see arrows??).

Later in development at stage 35, dorsal expression becomes restricted to rhombic lip of the fourth ventricle (Fig. 5A) and at least a subset of its migratory derivates (Wingate, 2001). In the hindbrain (Fig. 4B), *UNC5H3* labelling was found in the presumptive vestibuloacoustic nuclei (nucleus laminaris (NL), angularis (NA) and magnocellularis (NM)), the inferior olive (IO). In the cerebellum, the rhombic lip gives rise to the cerebellar granule cell precursor population, which forms the *Unc5h3*-positive external granule cell layer (EGL) covering the surface of the cerebellum at HH35/E10 (Fig. 5C). At E10, in transverse section, *UNC5H3*-positive granule cells from the EGL commence an inward, radial migration that is characterised, in the avian cerebellum, by the formation of parasagittal „raphes” (r) where cell density is highest (Fig. 5D). Again, this later pattern of embryonic *UNC5H3* is associated with the most dorsally derived neural tube populations at or near the roof plate interface.

**Figure 5.**
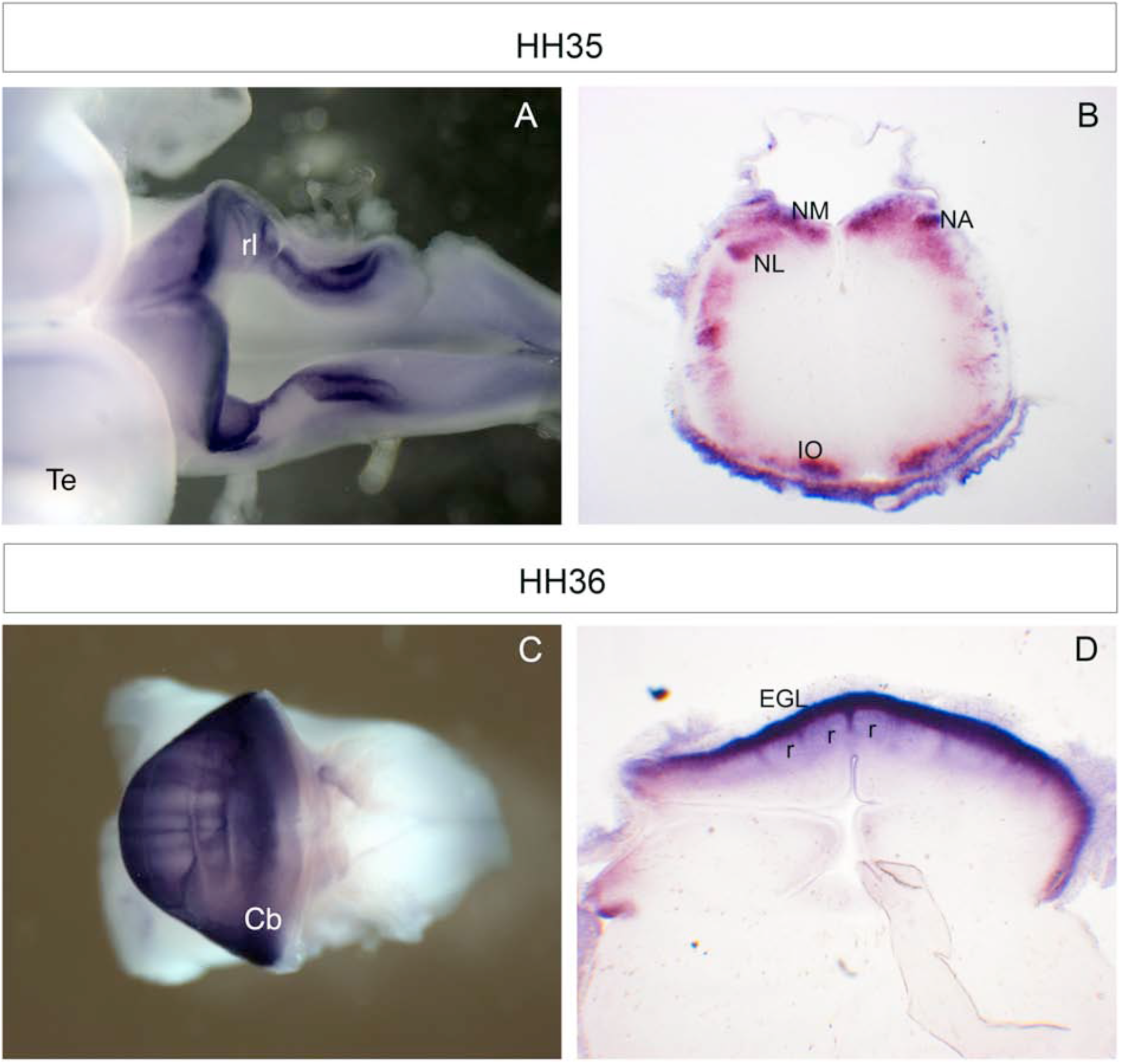
Whole-mount *in situ* hybridisation and transversal section of the rhombencephalon showing *UNC5H3* expression at later stages. In (A) and (C) anterior is left, posterior right; in (B) and (D) dorsal is up. (A) At embryonic day (E) 9 expression of *UNC5H3* becomes restricted to the rhombic lip. (B) In the hindbrain at E9 *UNC5H3* is expressed in the nuclei laminaris, angularis and magnocellularis, and in the inferor olive. (C) At E10 *UNC5H3* expression is detected in the external granule cell layer covering the whole surface of the cerebellum. (D) In transverse section of E10 cerebellum *UNC5H3* expression is seen in the EGL and raphes. Abbr.; Cb – cerebellum, EGL – external granular layer, IO – inferior olive, NA - nucleus angularis, NL – nucleus laminaris, NM – nucleus magnocellularis, r – raphes nuclei, rl – rhombic lib.

In summary, *UNC5H3* is in addition to its expression during cerebellar development, strongly expressed in motoneurones in the spinal cord, and the roof plate during a long period of time in the developing chick embryo. This may implicate its involvement in establishing or maintaining the identity of neuronal populations at the dorsal apex. In mouse, a mutation in Unc5h3 results in cerebellar granule cells migrating from rostal hindbrain into the midbrain (rostral cerebellar malformation; Ackermann et al., 1997), presumably through a failure in normal mechanisms of repulsion at the isthmus (Przyborski et al., 1998). A binding of GPC3 to Unc5 in revealed a role of Unc5 in guiding migrating pyramidal neurones in mouse cortex (Akkermans et al., 2021). Unc5 receptors can also act as dependence receptors, which can induce apoptosis without ligand (Llambi et al., 2001; Mehlen and Mazelin, 2003). Thus, neural cell death could happen without Netrin binding. Roof plate cells act as dorsal organiser cells producing BMP and WNT ligands that are necessary for the development of dorsal interneurons (Augsburger et al., 1999; Chizhikov and Millen, 2005; Gupta et al., 2022; Lee et al., 2000; Phan et al., 2010). Rekler et al (2024) unraveled *UNC5C/UNC5h3* expression in the roof plate as a novelty (Rekler et al., 2024). Our staining results confirm that UNC5c/UNC5H3 is expressed til at least HH 36 in the dorsal neural tube of chick embryos.

As the *NETRIN-1* knock-out mice lack an *UNC5H3* like phenotype, it was already speculated by Goldowitz et al. (2000) that the loss of NETRIN-1 may be compensated by an to date undiscovered Netrin familiy member or ligand that also binds to UNC5H*3*.

## 2. Experimental procedures

### Eggs

Fertile hen’s eggs were stored at room temperature and incubated on their sides in a humidified atmosphere at 37°C until they reached the desired stage. Embryos were staged according to Hamburger and Hamilton (1951). Embryos were fixed in 4% PFA and some embedded for the vibratom and sectioned at 50μm or in paraffin for *in situ* hybridisations.

### In situ hybridisation

W*hole-mount* mRNA *in situ* hybridisation was performed as previously described (Henrique et al., 1995). For the radioactive in situ hybridisation on paraffin sections, embryos were fixed in 4% paraformaldehyd, paraffin-embedded and sectioned at 8μm on a microtom. The radioactive *in situ* hybridisation was performed according to a modified version of Dagerlind et al. (1992).

The plasmid (ChEST 262h10) used for transcription of antisense probe, homologous to a 500 bp fragment near the 3’ end of gallus gallus *UNC5H3* mRNA, was obtained from the UK chicken EST Consortium. The plasmids containing fragments of *NETRIN 1*,and *NETRIN 2*, were a gift from M. Tessier-Lavigne and previously descibed in Serafini et al. (1994). The plasmid containing the *ISLET 1* sequence was a gift from Thomas Jessel.

### Whole-mount immunohistochemistry

Following fixation in 4% PFA embryos were washed in PBS containing 0,1% Triton X-100 and 10% fetal calf serum in and then incubated for 2-3 days at 4°C with anti-Islet1/2 antibody (Developmental Studies Hybridoma Bank; 1:1 in PBS, 0,1% Triton X-100, 10% FCS). Then, the embryos were washed several times and incubated with a fluorchrome-conjugated antibody (Cy3-anti-mouse; 1:200; Dianova, Hamburg, Germany) for 1 days at 4°C. Embryos were given several final washes before embedding for vibratom sections.

